# Socially learned call sequences reveal gradual development of higher-order structure

**DOI:** 10.64898/2026.06.15.732496

**Authors:** Stephanie L. Mason, Sarah L. Walsh, Stephanie L. King, Amanda R. Ridley

## Abstract

Syntax was long considered to distinguish human language from other vocal systems, with parallels in non-human animals historically limited to song. However, song lacks discrete meaning, which is a crucial pre-requisite of linguistic syntax. Over the last two decades evidence of combinatoriality in the discrete, semantic calls of an array of taxa has accumulated, providing the opportunity to investigate potentially closer parallels to language. However, most examples remain limited to small repertoires of simple two-call sequences, preventing evidence of complex internal structuring like that seen in human sentences. The recent discovery that several species produce extensive repertoires of much longer call sequences, has provided the opportunity to investigate the full extent of syntactic structure in non-human call systems. Here we demonstrate that Western Australian magpies (*Gymnorhina tibicen dorsalis*) use multi-level structured ordering rules within their semantic call sequences and that these ordering rules are learned during development. Specifically, we find that calls within sequences up to 15 calls long depend on the two calls given prior and that independently produced segments (‘phonemes’), calls, and sequences recombine into longer structures, indicating hierarchical organisation. This represents the first evidence of multi-level non-adjacent organisation and learned syntactic structure in a semantic non-human system.

## Introduction

At the core of language lies syntax—the mechanism by which we combine otherwise meaningless sounds into meaningful words (*phonology* or *phonological syntax*), and those words into an infinite array of structured, generative phrases (*compositional syntax*) that enable the extraordinary breadth and flexibility of human communication (Hurford, 2012; Marler, 1998). Given the importance of syntax to linguistic complexity, and the fact that its evolutionary origins remain unresolved (Christiansen & Kirby, 2003; Hauser & Fitch, 2003; Weiss & Newport, 2006), it is unsurprising that considerable research has sought to uncover comparable structures in non-human vocal systems (Berthet et al., 2025; Berwick et al., 2011; Bortolato et al., 2023; Engesser et al., 2016, 2019; Engesser & Townsend, 2019; Girard-Buttoz, Zaccarella, et al., 2022; Hedwig & Kohlberg, 2024; Leroux & Townsend, 2020, 2020; Selbmann et al., 2023; Suzuki, 2014; Suzuki et al., 2017, 2020; Walsh et al., 2023).

Traditionally, syntactic research in non-human animals has focused on bird and whale song, in which meaningless syllables are combined into complex songs that are holistically functional: their quality conveying individual quality or a threat to rivals (Hurford, 2012; Marler & Pickert, 1984). Song can be structurally complex, sometimes showing complex hierarchical structural organisation (Berwick et al., 2011; Doupe & Kuhl, 1999; Katahira et al., 2011; Menyhart et al., 2015; Payne & McVay, 1971; Sainburg et al., 2019; Soma et al., 2009; Tchernichovski et al., 2001), but it lacks semantic meaning—meaning may be inferred from the overall quality of the song but it cannot be broken down into discrete meaningful units. As such, song can only be compared to phonological syntax, with its hierarchical *structural* levels operating exclusively within this single *combinatorial/syntactic* level. While song research has been invaluable to our understanding of complex cognitive processes like vocal learning and structural complexity (Arnon et al., 2025; Audet et al., 2023; Beecher & Akçay, 2020; Doupe & Kuhl, 1999; Marler, 1970; Marler & Pickert, 1984; Menyhart et al., 2015), its lack of semantic meaning limits meaningful comparison with linguistic syntax, for which semanticity is a crucial pre-requisite (Berwick et al., 2011; Bolhuis et al., 2010; Engesser & Townsend, 2019). More recently, a shift to studying the semantic, non-song (i.e. call) repertoires of various taxa has revealed the presence of syntactic-like structure akin to both phonological (e.g. chestnut-crowned babblers, *Pomatostomus ruficeps*, Engesser et al., 2019) and compositional syntax (e.g. Japanese tits, *Parus minor*, Suzuki et al, 2017; bonobos, *Pan paniscus*, Berthet et al., 2025). Evidence of syntactic-like structure in vocalisations that possess discrete, context-specific meaning presents promising opportunities for comparative research into the possible evolutionary origins and drivers of this capacity outside of human language.

Most studies on semantic combinatoriality have found only two-call combinations, and combinatoriality that parallels only one of the levels seen in linguistic syntax; i.e. either phonological combining of meaningless sounds into meaningful calls (Engesser et al., 2019; Ouattara et al., 2009) *or* combining meaningful calls into sequences (Arnold & Zuberbühler, 2006; Hedwig & Kohlberg, 2024; Collier et al., 2020; Collier et al., 2017; Déaux et al., 2016)—with a small number showing evidence of compositional rules at this level (Suzuki et al., 2016, 2018, 2020; Berthet et al., 2025)—but not both (though see Walsh et al., 2023 for the first evidence of multi-level combinatoriality). This may be because there is a genuine discontinuity between these vocal systems and linguistic syntax, or may be the result of almost all research into call combinatoriality focussing on either a single or small number of vocalisations in the species’ repertoire (e.g., Engesser et al., 2016, 2019; Hedwig & Kohlberg, 2024; Ouattara et al., 2009; Suzuki et al., 2016; Walsh et al., 2019). While the lack of whole-repertoire studies is likely partly because analysing structure in even a few short vocal sequences is incredibly time-consuming, it has potentially limited our capacity to uncover the true extent of syntactic capacity within these vocal systems (Berthet et al., 2025; Bosshard et al., 2024; Girard-Buttoz, Zaccarella, et al., 2022; Walsh et al., 2023). Linguistic syntax is undoubtedly unparalleled in flexibility and generative capacity, but whole-repertoire investigations are needed to fully capture the extent of parallels in other animals.

Four recent studies have attempted to address this gap using whole repertoire approaches, and in doing so, have found evidence of some of the most complex syntactic capacity shown in animal systems to date (Berthet et al., 2025; Bosshard et al., 2024; Girard-Buttoz, Zaccarella, et al., 2022; Walsh et al., 2023). Girard-Buttoz et al. (2022) found that chimpanzees (*Pan troglodytes*) combine calls into predictable pairs (bigrams) and triplets (trigrams), where preference for certain bigrams is retained within trigrams (e.g. A-B appears in the trigrams A-B-C and A-B-D, maintaining its first-order or pairwise structure), suggesting positional regularities and recombination are present in chimpanzee sequences. The study recorded 390 unique sequences, ranging in length from two to 10 units long (Girard-Buttoz, Zaccarella, et al., 2022)—the most extensive known use of non-song combinatoriality in any animal system. Utilising Markov chain models, Bosshard et al. (2024) was able to show the presence of second-order structure—where the probability of a given call being produced in sequence relied on not only the previous call, but the one before that—and some amount of recombination in the sequences of common marmosets (*Callithrix jacchus*), which can be up to 9 units long. The cooccurrence of second-order structure and recombination is suggestive of hierarchy and represents some of the first evidence of such structuring in a non-song animal vocal system, highlighting the need for further investigation into the potential for similar structure in other species. While similarly long sequences have yet to be shown in bonobos (*Pan paniscus*), Berthet et al. (2025) was able to show that three bonobo bigrams show evidence of non-trivial compositionality—that is, the meaning of A-B is derived from but distinct to the meanings of A and B individually, but also from the simple sum of their meanings (i.e. A followed by B). While less structurally complex than the evidence in marmoset sequences, the presence of three distinct, non-trivial call combinations represents a first in any non-human system (though see Suzuki et al., 2016 for evidence in a single call combination).

Given non-human primates appear to lack the capacity for vocal production learning—the ability to learn new vocalisations from heard models—syntax has been proposed to have developed within the primate lineage to mitigate the communicative constraints of a genetically fixed repertoire (Bortolato et al., 2023; Nowak et al., 2000), suggesting humans evolved the capacity for vocal production learning after our divergence from other primates. Western Australian magpies, like humans, are life-long vocal production learners—not only can they learn to produce novel vocalisations, as evidenced by their capacity for mimicry, but they can do so across their entire lifespan (Kaplan, 1999; Suthers et al., 2011). Alongside their song and mimicry vocalisations, magpies produce context-specific, meaningful calls (Dutour et al., 2020, 2023; Walsh, 2024) that they combine into over 100 unique sequences that range from 2 to 15 calls long (Mason 2025; Mason et al., 2026; Walsh, 2024; Walsh et al., 2019, 2023, 2024). While it is infeasible to determine the meaning of every individual call and sequence in a system so diverse, preliminary evidence confirms that individual calls and sequences are used in different contexts, predominantly relating to threats (Mason, 2025; Walsh, 2024; Walsh et al., 2019). One clear example is a general disturbance alarm call that elicits low-level vigilance and horizontal scanning, that combined with a two-call alert sequence, forms a three-call aerial threat sequence that elicits heightened vigilance and upward scanning (Mason, 2025). Notably, magpies are the first species shown to combine at both the within-call (phonological) and between-call levels (Walsh et al., 2019, 2023; Walsh, 2024). Initial findings by Walsh et al, (2023) reveal that magpies exhibit at least first-order dependencies both between ‘segments’ (akin to phonemes) within calls, and calls within sequences—but longer-range dependencies are yet to be investigated. The presence of two key features thought to set human language apart from other vocal systems in complexity—open-ended vocal production learning and extensive combinatoriality—paired with being the first species shown to use multi-level ordering rules within semantic call sequences, makes magpies a particularly promising model system to explore alternative theories for why syntax may have evolved.

To this end, our recent research documented the ontogeny of these vocalisations in fledgling magpies and showed that the call sequence repertoire is learned from the social group, with more sociable fledglings learning faster and developing larger repertoires (Mason et al., 2026). Our findings suggested a process where fledglings first produce segments, quickly followed by simple two-segment calls (formed through phonological combining), before learning to combine calls into group-specific sequences by listening to the call sequences of social contacts (Mason, 2025; Mason et al., 2026). While learned structuring of songs is well studied and often compared to language learning (Berwick et al., 2011; Bolhuis et al., 2010) an important distinction is the lack of meaning associated with song (Suzuki et al., 2018). Our findings represent the first evidence of learned semantic call sequences in any non-human animal (Mason et al., 2026), mirroring, to some extent, the process of language learning in infants: phonemes combined into meaningful calls/words are first produced in isolation, before being combined into increasingly longer sequences/sentences learned through social exposure (Yang, 2013; Yang et al., 2017). While magpies are open-ended vocal production learners, acoustic analyses revealed that segments do not undergo acoustic development—a prerequisite for evidencing vocal production learning (Egnor & Hauser, 2004)—suggesting the segment ‘building blocks’ of calls and sequences are innate (Mason, 2025). Now, structural analysis is needed to confirm that it is instead the ordering rules within sequences that are being learned through a process more akin to usage learning—i.e. learning the correct way in which to combine calls. Inspired by their use in Bosshard et al. (2024) and building upon the findings of Walsh et al. (2023), we used Markov chain models to further explore the extent of structure in magpie call sequences (i.e. beyond the first order) and investigate differences in structure between age groups. Specifically, we examine whether first- or second-order Markov models best capture the structure of magpie call sequences and whether transitions used by adults and fledglings differ at the first- and second-order levels.

## Methods

### Study species and population

Western Australian magpies are passerines that live in stable, cooperatively breeding social groups (Pike et al., 2019). They can live up to 30 years, reaching sexual maturity at between 1-3 years of age when their sex-specific plumage develops (Pike et al., 2019; Rowley et al., 2022). Magpies are open-ended vocal production learners that produce a range of song and non-song vocalisations including mimicry, choruses, carolling, discrete calls and call sequences (Brown & Farabaugh, 1991; Dutour et al., 2020; Kaplan 1999; Walsh et al., 2019, 2023). Within their discrete call and call sequence repertoire, magpies produce four sound segments: ‘noisy line’ (*NL*), ‘down-sweep’ (*DS*), short-high (*SH*) and long-high (*LH*). These segments may be produced independently as stand-alone, single-segment calls—specifically *DS*, *NL* and *LH* (SH is not produced in isolation), with meaning identified in at least *NL* (i.e. *NL* functions as a general disturbance alarm call, Mason, 2025; Walsh, 2024)—or combined into multi-segment calls: predominantly *NLDS, SHDS* and *LHDS* (see Walsh et al. 2023 for evidence of longer calls). The segments within these multi-segment calls are delineated only by sudden spectral shifts and are produced as one continuous sound. Next, these single- and multi-segment calls are combined into a vast number of different sequences ranging from 2 to 15 calls long (with gaps of ∼0.1 to <0.5 seconds between each call) (Figure 1).

**Figure 1.**
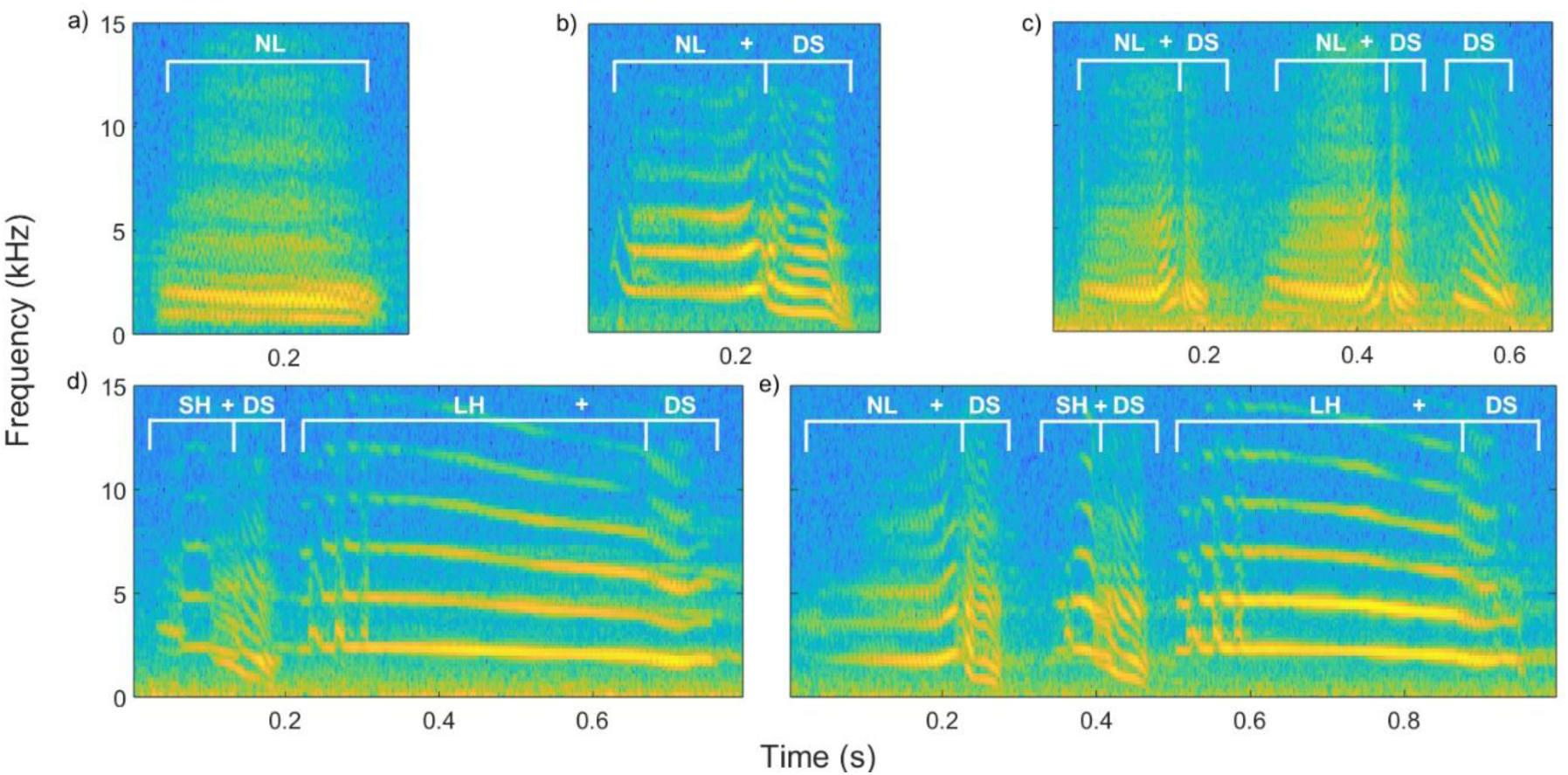
Examples of magpie discrete calls and call sequences. Vocalisations shown were recorded at 44.1kHz sample rate and spectrograms were generated with a 256-point FFT with a Hann window and 250-sample overlap leading to a frequency resolution of ∼172.3Hz. Labels indicate distinct call segments: ‘*down sweep’* (*DS*), ‘*long high’* (*LH*), ‘*noisy line’* (*NL*) and ‘*short high’* (*SH*), which can be produced in isolation (single-segment calls) as in *NL* (a) or combined into multi-segment calls as in *NLDS* (b). Segments within the same multi-segment call are produced as one continuous sound and are delineated only by sudden spectral shifts (shown here under the same bracket with segment labels separated by ‘+’). Calls within call sequences (c, d, and e) are separated by brief silences (< 0.5s). Consecutive repetitions (e.g. the 2 repetitions of the *NLDS* call in (c)) were not removed for analysis to directly address the extent of redundancy in this vocal system. Refer to Walsh et al., 2023 for further detail on magpie vocal classification.

Data for this study was collected by SLM from eight social groups situated in the Perth metropolitan areas of Crawley (31.97°S, 115.82°E; n = 5 groups) and Guildford (31.89°S, 115.97°E; n = 3 groups) ranging in size from 2 to 9 adult members, with between 1-6 fledglings present in any one group at a time. The groups are part of a long-term study population (since 2014) and are habituated to human presence, allowing for natural observation and recordings from <10m (Speechley et al., 2024; Walsh et al., 2023). Individuals are identifiable either by coloured rings or by distinctive physical features (e.g., plumage variation or scarring). Fledglings are distinguishable from adults by their lack of sexually mature plumage: fledglings have mottled grey plumage, whereas adults are black—adult males have white backs and adult females have black and white mottled backs.

### Data collection

Vocal data was collected during focal follows conducted as part of a broader study (Mason, 2025; Mason et al., 2026) on fledgling magpies and their social groups (n = 11 fledglings from 11 mothers in 8 social groups) from October 2022 to June 2023. One-hour focal follows were conducted from the first week to 200 days post-fledging, once a week for the first 100 days, and then every 3 weeks to 200 days. A total of 196 focal follows were conducted, amounting to ∼145h of observation time and recording. During follows, continuous audio recording captured all vocal productions of focal fledglings and their group members, along with accompanying voice notes identifying the caller (Mason et al., 2026).

### Data preparation

Audio files were examined spectrographically in Adobe Audition (v22.3) and individual vocal events were manually identified and classified using established methods (*sensu* Walsh et al., 2023, 2024; see Mason, 2025 for more detail). Only call sequences were analysed in the current study, which are sequences of different calls uttered with a maximum of 0.5s between each constituent call type, where individual vocal events produced by a caller are delineated by silences of >0.5s (Walsh et al., 2023). While consecutive repetitions of calls within sequences are often removed due to them being unlikely to encode new meaning (e.g., A-A-B-C would become A-B-C, Girard-Buttoz, Zaccarella, et al., 2022; Mason et al., 2026; Walsh et al., 2023), this may ignore the structural significance of redundancy patterns. In human language, consecutive repetition of units is rare and typically constrained to specific morphological or phonological contexts (e.g., emphatic repetition: “very very bad”; reduplication: “so-so”; baby-talk forms: “mama” and “papa”; Bazzanella 2011). Removing such repetitions from analysis may therefore overestimate the system’s structural similarity to linguistic syntax by obscuring a key difference in how information is encoded. Given that in magpies there is evidence that repetitions encode urgency rather than distinct meaning (Dutour et al., 2023; Walsh et al., 2023, 2024), for the purpose of assessing the extent of internal structuring we did not remove consecutive repetitions and instead directly addressed their presence. Where an entire sequence was repeated within a single vocalisation, this was reduced to a single iteration (*NLDS-SHDS-LHDS-NLDS-SHDS-LHDS* became *NLDS-SHDS-LHDS*). We believe this is a different case to consecutive repetition of calls, and likely functions in salience in high urgency situations—i.e. we would not say ““we we need need to to go go!” but might say “we need to go! we need to go!” to ensure successful communication. Note that some primate studies remove any repeated units consecutive or otherwise—i.e. A-B-A-C becomes A-B-C—but this can be considered overly conservative as even in human language we may repeat the same word in a sentence to generate additional meaning (e.g. “<u>you</u> said <u>you</u> would call!”) (Girard-Buttoz et al. 2022). The call composition of each sequence was extracted along with caller identity.

For each caller, unique sequence types were identified and the number of times that they produced each type was tallied. This meant each sequence type was only listed once per individual but could be listed across multiple individuals, accompanied by individual frequency values. Each caller was coded as either a fledgling (0) or adult (1). We excluded non-focal fledgling vocalisations. Sequences were converted into long format such that each row of the CSV data file contained one constituent call, with the following row containing the call that followed it within the sequence. The first and last row for each sequence contained ‘Start’ and ‘Stop’ respectively, so that the probability of calls and pairs of calls starting or ending a sequence could be determined.

Our final sample size was 1892 sequence productions (fledgling n = 548, adult n = 1344). The following statistics aim to describe the diversity of call sequences and as such, ignore consecutive repetitions to provide conservative estimates (though note these repetitions were not excluded during analysis). The data set comprised 108 unique sequence types (see Mason et al., 2026 for the full 119 sequence types before non-focal fledgling exclusion), varying in length from 2 to 15 calls long (excluding ‘Start’ and ‘Stop’ states). 38 of these sequence types were produced by both adults and fledglings, 36 only by adults, and 34 only by fledglings (fledgling n = 72 types, adult n = 74 types). Two-call sequences comprised 57.7 and 52.7% of the fledgling and adult sequence productions respectively, three-call sequences were 26.8% and 39.8%, four-call sequences were 10.8% and 6.43% respectively and call sequences ≥ five calls were 4.75% and 1.06% respectively. Total frequency of sequence types ranged from *NLDS-SHDS-LHDS* and *SHDS-LHDS* that were produced 356 and 453 times respectively across the study by both adults and fledglings—to sequences produced only once (x̅ = 18.25 ± 6.18). Rather than exclude these rare sequences that may represent important variability or flexibility in fledgling development, we included frequency weighting of transitions in the analysis (see below).

### Markov chain models

To assess the presence of internal structure in magpie sequences, they were modelled using Markov chain models. Specifically, we developed custom MATLAB scripts that calculated first-order Markov transition probabilities (the probability of call B occurring after call A) and second-order Markov transition probabilities (the probability of call C occurring given A → B has already occurred). First-order Markov chains assume that only the current state predicts the subsequent state, whereas second-order chains assume that both the current and previous state predict the subsequent state. While second-order dependencies may not capture the presence of potential longer-range sequencing, analysis beyond the second order would have required reducing the dataset considerably with each additional order, to remove sequences of insufficient length. Nonetheless, the presence of second-order structure is sufficient to demonstrate a level of call organisation that matches the highest complexity reported in any non-human semantic call system to date (i.e. common marmosets, Bosshard et al., 2024).

Every consecutive pair and triplet of calls within each sequence was extracted from the fledgling and adult datasets. Note that ‘bigram’ and ‘trigram’ refer to sequences of exactly 2 and 3 calls long respectively, so ‘pairs’ and ‘triplets’ are used to also capture those cut from within longer sequences. For example, the sequence *NLDS-SHDS-LHDS* would yield the pairs: *Start-NLDS*, *NLDS-SHDS*, *SHDS-LHDS* and *LHDS-Stop* and the triplets: *Start-NLDS-SHDS, NLDS-SHDS-LHDS* and *SHDS-LHDS-Stop*. Transition counts were weighted by the total frequency with which they appear in the dataset, including when they appear multiple times within a sequence (e.g. the pair *NLDS-SHDS* occurs twice in *NLDS-SHDS-NL-NLDS-SHDS*), and when that sequence is produced multiple times (both within and across individuals). First order transition probabilities were calculated by the following formula:

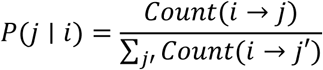

Where the probability of transitioning to call *j* given call *i* is the number of times *j* did follow *i*, divided by the total count of *i* transitioning into any call. This normalises transitions to the sum of 1 in each row such that the total probability of call *i* transitioning to any other call, or to ‘stop’, is equal to 1 (i.e. total probability for that call).

Second order transition probabilities were calculated using the formula:

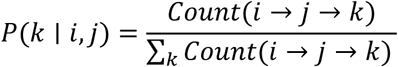

In which the number of times call *k* occurred after transition *i* → *j* is divided by the total number of times any call followed the transition *i* → *j*.

### Statistical analysis

All statistical analysis was performed using custom MATLAB scripts. While the frequency weighted transitional probabilities (tP) tell us how likely a particular transition is within the observed data, this only describes the data and does not necessarily indicate the presence of meaningful structure. If A → B occurs 40% of the time for example, this may still be what we would expect by chance based on the frequency of these calls in the dataset. To assess the significance of apparent structure, transition probabilities must be compared to a null model of what would happen if there was no structure—i.e. if calls were combined randomly—to determine if there is a deviation from expectation. As such, we performed 10,000 permutations of randomly reconstructing sequences using the base frequencies of each call in the dataset, preserving the position of *Start* and *Stop* and the empirical distribution of sequence lengths (i.e. all randomly generated sequences were of realistic length and capped at 15 calls as per the observed maximum, with individual calls appearing in the same frequencies as the observed data). Transition probabilities were recomputed for each permutation of the data to form a null distribution of “expected” chance probabilities. This was done separately for the adult and fledgling data, for both their first and second order transitions. The observed transition probabilities were then compared to the null distribution of expected probabilities using two-tailed permutation tests to test whether the observed values differed from chance (with an alpha level of 0.05) after accounting for baseline call frequencies and sequence lengths. Two-tail tests were performed as observed frequencies below what would be expected by chance are just as indicative of structure as observed frequencies above what would be expected by chance—the former points to avoided transitions and the latter to preferred transitions. To control for multiple comparisons across the transition matrix, p-values were corrected using the Benjamini-Hochberg false discovery rate (FDR) procedure (Benjamini & Hochberg, 1995), applied separately for each group (adults and fledglings) and each Markov order. Transitions with FDR-corrected p-values below 0.05 were considered significant. To identify whether significance was indeed due to significantly lower or higher probability than chance, the randomly permuted probabilities were averaged to form expected probability matrices for comparison to the observed matrices. This allowed us to conclude the presence of real structure at the individual sequence level.

### Model comparison

To assess whether the first- or second-order model explained the adult and fledgling repertoires best, total log-likelihood (loglik) values calculated across all frequency-weighted transitions in the adult and fledgling data respectively were compared. Loglik values were computed by summing the log probabilities of each observed transition under the respective model. More positive (less negative) loglik values indicate better model fit. To handle zero probabilities associated with transitions that never occurred, Dirichlet (additive) smoothing was applied, adding a small constant k to all transition counts prior to normalization (Jurafsky & Martin, 2025). Six values of the smoothing parameter (*k* = 0.0001, 0.001, 0.01, 0.1, 1, 10) were tested to assess the robustness of the results to smoothing strength (Bosshard et al., 2024). We found that k=10 over-smoothed slightly and caused underfitting of the data, but that loglik values were constant for k = 0.0001-1 suggesting the presence of robust and consistent sequential structure. As such, results are presented averaged over k = 0.0001-1 (see electronic supplementary material Figures S.1. and S.2. for full comparison of k values). The smoothed first- and second-order transition probability matrices were compared to a uniform, zero-order transition matrix (where every possible transition is assigned equal probability) that assumes calls are combined randomly, by comparing total loglik values across all models and k-values. While adult and fledgling models cannot be directly compared due to differences in baseline frequencies, sequence length and observed transitions, loglik values were normalised to account for differences in dataset size. This allowed us to compute average loglik per-transition, placing the adult and fledgling loglik values on the same scale for relative comparison (though they should not be interpreted as absolute measures).

## Results

### First-order Markov chains

#### Adults

P-values reported throughout were calculated using two-tailed permutation tests, as described in Methods. At the first-order level, clear structure is apparent, with a total of 11 transitions produced significantly more than chance expectation and 17 transitions produced significantly below chance expectation (Figure 2a). *NLDS* was the only call with an above-chance probability of starting a sequence (*p* < 0.0001), and was the only call to significantly transition into *DS*, though probability was relatively low (tP = 0.16, p < 0.01).

**Figure 2.**
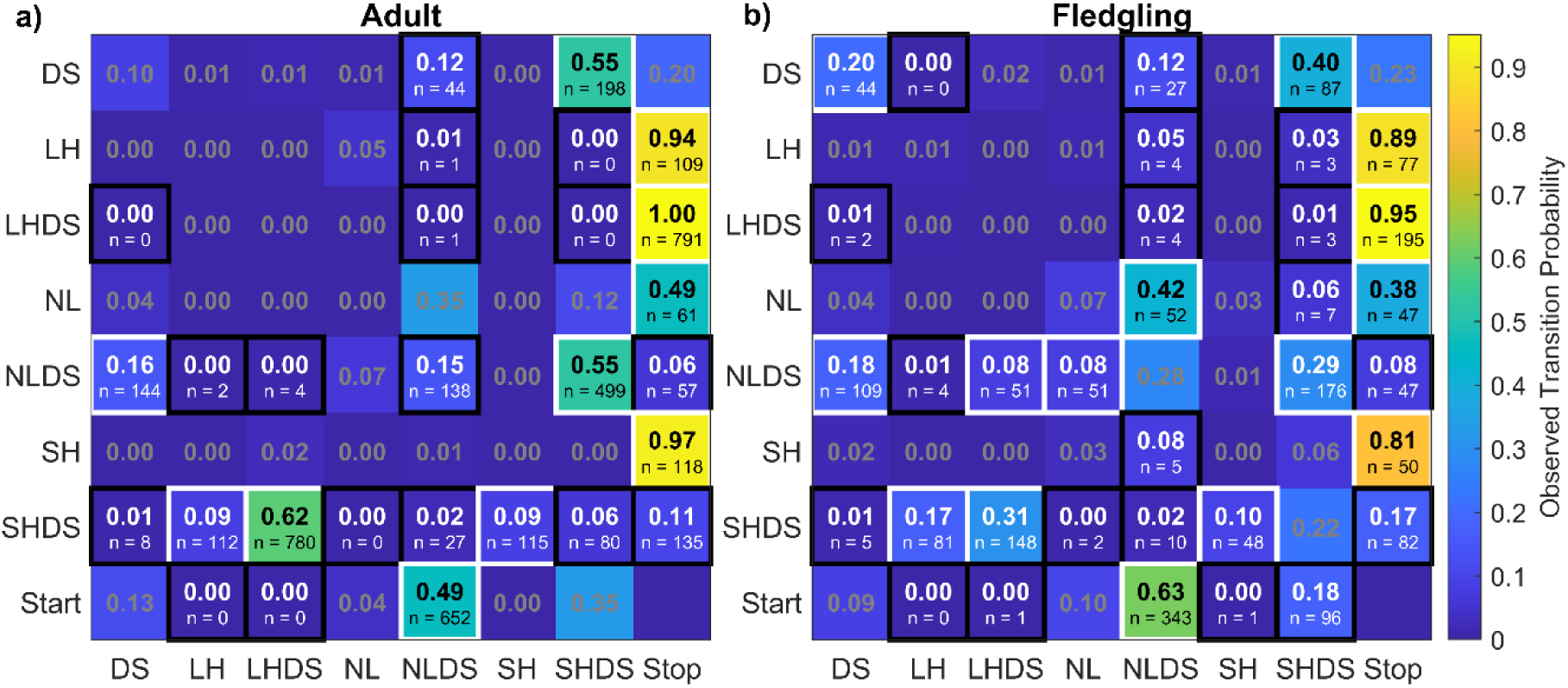
First-order Markov chain model matrices depicting for (a) adult and (b) fledgling magpies, the transition probabilities from a call or *Start* state (rows) to the next call or *Stop* state (columns). The diagonal values from top left to bottom right show same-to-same call transition probabilities. Values in cells are the observed probabilities—more yellow cells for higher probabilities and more purple cells for lower probabilities. Cells with a white border show values that were significantly higher than chance expectation, while those with a black border show those that were significantly lower than chance expectation, when compared to a null distribution of 10,000 permutations of randomly generated sequences using the baseline frequency of calls and distribution of sequence lengths in the data. Sample sizes for significant transitions are shown below the probabilities. *Start* → *Stop* transitions are not possible so are also blank.

*LH, LHDS*, *NL* and *SH* function as terminal calls, transitioning exclusively to Stop above chance (*LH, LHDS, SH* all *p* < 0.0001, *NL p* < 0.01), suggesting they are avoided mid-sequence. *DS* and *NLDS* both transitioned preferentially to *SHDS* (*p* < 0.0001), while *SHDS* tended to transition to *LH*, *LHDS*, and *SH* (all *p* < 0.0001), though *SHDS-LH* and *SHDS-SH* probabilities were relatively low (both tP = 0.09).

The first order matrix shows low diagonality: only *DS, NLDS* and *SHDS* were ever repeated consecutively. *DS-DS* occurred at chance levels (tP = 0.10), while *NLDS-NLDS* and *SHDS-SHDS* were actively avoided (*p* = 0.013 and *p* < 0.0001 respectively). Several calls were avoided at the start of sequences (*LH*: *p* < 0.01 and *LHDS*: *p* < 0.0001) and at the end of sequences (*NLDS, SHDS*: *p* < 0.0001). Numerous additional transitions occurred significantly below chance (indicated by black borders in Figure 2a; p-values given in Supplementary Table S.1.). The presence of many significantly below-chance transitions is suggestive of intentionally structured sequences.

#### Fledglings

Fledglings show similar first-order structure to adults (Figure 2b), with all 11 above-chance adult transitions also occurring above chance in fledglings. However, fledglings produce four additional above-chance transitions not seen in adults: they show significant repetition in *DS-DS* (*p* < 0.0001), which occurred only at chance in adults; they transition from *NL* to *NLDS* above chance (*p* < 0.01) which was also occurred relatively frequently (tP = 0.35) but not above chance in adults; and transition from *NLDS* into *LHDS* and *NL*, though both with low probability (tP = 0.08; *p* = 0.025 and *p* = 0.026 respectively). In total, fledglings produced 15 transitions significantly above chance.

Fledglings avoid 18 first-order transitions, largely overlapping with the 17 avoided by adults, but with notable differences. Unlike adults, fledglings do not avoid same-call repetitions: *NLDS-NLDS* and *SHDS-SHDS* occur at chance levels, while *DS-DS* occurs above chance (see above). Fledglings avoid additional calls at the start of sequences (*Start-SH*: *p* = 0.012; *Start-SHDS*: *p* < 0.001) that adults do not and avoid *NL-SHDS* (*p* < 0.01), which occurs at chance levels in adults. Several other minor differences exist: *DS-LH* and *SH-NLDS* are avoided (*p* = 0.044 and *p* = 0.025 respectively) which occur at chance levels in adults and *NLDS-LHDS* occurs above rather than below chance in fledglings (see Figure 2b for full transition matrix and Table S.1. for p-values).

### Second-order Markov chains

#### Adults

At the second-order level, there is clear evidence of structured call transitions (Figure 3a; Supplementary table S.2.). The most frequent transitions—*Start-SHDS-LHDS, SHDS-LHDS-Stop, Start-NLDS-SHDS,* and *NLDS-SHDS-LHDS*—show the presence of overlapping significant transitions that reflect two common sequences: *‘SHDS-LHDS’* (associated with recruitment; Walsh, 2024) and ‘*NLDS-SHDS-LHDS’* (associated with aerial predator threats; Mason, 2025)—the latter representing recombination of a general disturbance alarm call (‘*NLDS’*) with the recruitment sequence. *Start-SHDS* exclusively transitions to *LHDS* above chance (*p* < 0.0001) and *SHDS-LHDS* exclusively transitions to *Stop* above chance (*p* < 0.01), indicating ‘*SHDS-LHDS’* is a highly stable call sequence. Similarly, *Start-NLDS* only transitions to *SHDS* above chance levels (*p* < 0.0001). *NLDS-SHDS* shows more variability, transitioning above chance to *LHDS* (most common, *p* < 0.0001), *LH*, or *SH* (*p* < 0.01 and *p* = 0.016 respectively, though with lower probabilities: tP = 0.15 and 0.11). Critically, all three variants—*SHDS-LHDS, SHDS-LH,* and *SHDS-SH*—transition almost exclusively to *Stop* (*SHDS-LH-Stop* and *SHDS-SH-Stop* are both highly probable tP = 0.95 and 0.99 respectively but are insignificant, likely due to the relative rarity of *LH* and *SH* in the dataset: n = 203 and n = 184 of 3681 respectively, or ∼5% each of calls produced within adult sequences). These combined constraints (*Start-SHDS* to *LHDS*, *Start-NLDS* to *SHDS*, *NLDS-SHDS* to {*LHDS, LH, SH*} and *SHDS-*{*LHDS, LH, SH*} to *Stop*) suggest transitional dependencies spanning up to five sequential positions, extending well beyond triplet-level patterns (Figure 5).

**Figure 3.**
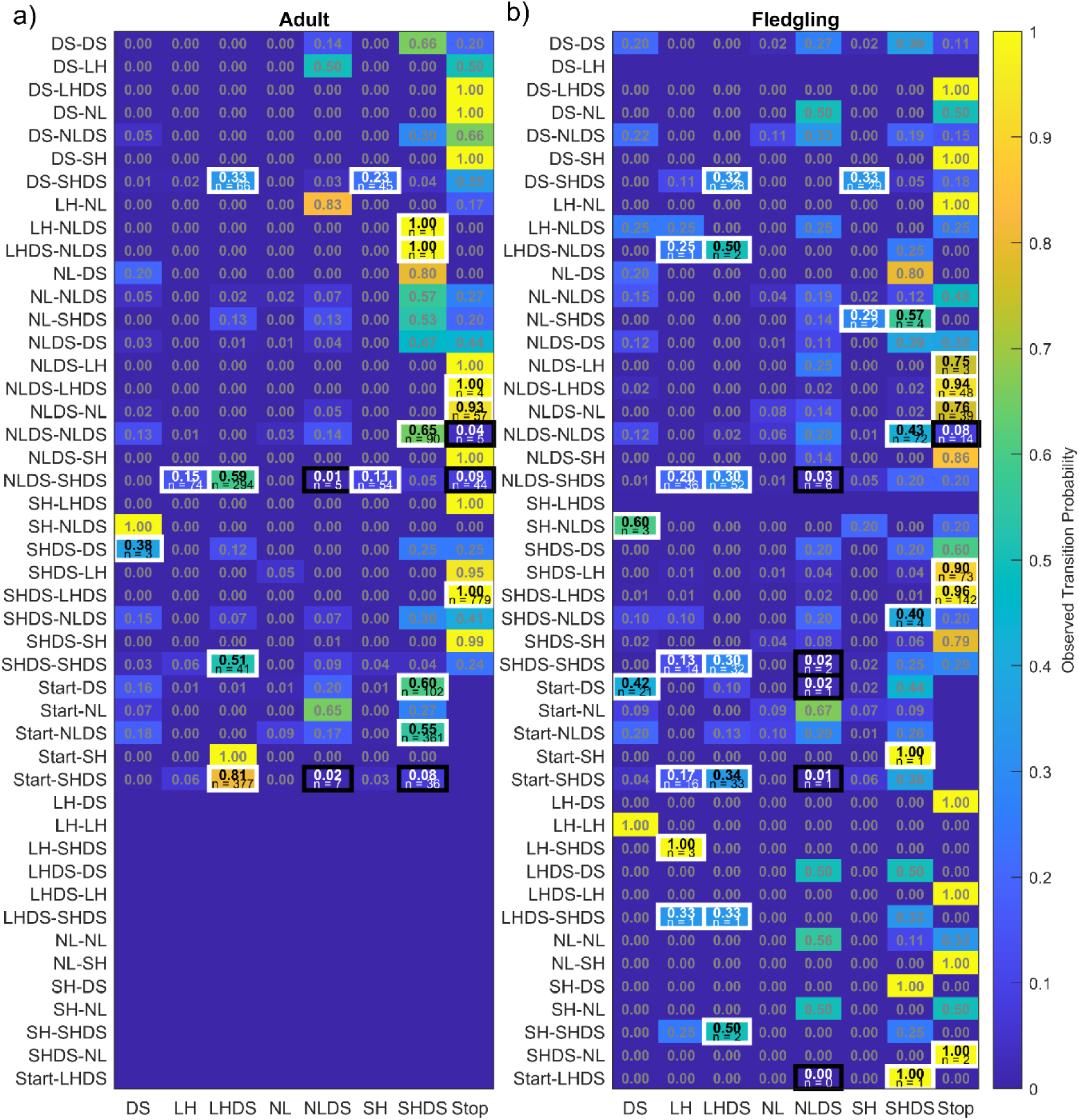
Second-order Markov chain model matrix depicting for (a) adult and (b) fledgling magpies, the transition probabilities from a pair of calls (rows) to a third call or *Stop* state (columns). Values in cells are the observed probabilities—yellow cells for higher and purple cells for lower probabilities. White borders indicate transitions that occurred significantly more than chance expectation and black borders, those that occurred significantly less than chance expectation when compared to a null distribution of 10,000 permutations of randomly generated sequences using the baseline frequency of calls and distribution of sequence lengths in the data. Sample sizes for significant transitions are shown below the probabilities. Blank rows indicate the absence of a transition in either the fledgling or adult dataset. *Start* → B → *Stop* transitions are not possible so are also blank.

When same-call repetitions occurred (which were generally avoided; Figure 2a), they showed structured transitions: *NLDS-NLDS* transitioned preferentially to *SHDS*, and *SHDS-SHDS* to *LHDS* (both p ≤ 0.0001). Rather than occurring randomly, repetition appears reserved for specific sequential contexts, suggesting functional constraint. Adults actively avoided 5 second-order transitions, including *NLDS-SHDS* to *NLDS* and *Stop*; *NLDS-NLDS* to *Stop*; and *Start-SHDS* to *NLDS* and *SHDS* (full results in Figure 3.a. with *p*-values given in Supplementary Table S.2.).

#### Fledglings

Fledglings show considerably more variable second-order structure than adults, with 13 additional transitions unique to fledglings versus only two unique to adults (represented by blank rows in the other age group’s figure: Figure 3b, Supplementary Table S.2.) and 28 transitions produced above chance levels compared to the 16 in adults. Like adults, fledglings produce the core terminal sequences *SHDS-LHDS-Stop* and *SHDS-LH-Stop* above chance, as well as *Start-SHDS-LHDS* and *NLDS-SHDS to LH* and *LHDS*. In contrast to adults, fledglings produce *Start-NLDS-SHDS* at chance levels—one of the most common and probable adult alarm sequences.

Unlike adults, fledglings seem to actively repeat calls, with *Start-DS* transitioning to *DS* significantly above chance (*p* < 0.0001). Fledglings produced numerous additional significant transitions at very low counts (n ≤ 5; see Figure 3a and Supplementary Table S.2. for complete list), including several terminal pairs (i.e. those transitioning to *Stop*) produced at chance levels in adults. These may represent exploratory vocal behaviour rather than functionally meaningful structure. In total, fledglings avoided 6 second-order transitions—3 of the same avoided transitions as adults (*NLDS-SHDS* to *NLDS*; *NLDS-NLDS* to *Stop;* and *Start-SHDS* to *NLDS*) as well as 3 fledgling-specific avoidances (full details in Figure 3b and Supplementary Table S.2.).

### Model comparison

While significant structure at both the first and second-order levels is shown above, this does not tell us if the second-order model is a better fit to explain the structure of the data. As such, the loglik of a random model—where the likelihood of each transition occurring is equal (0^th^ order)—was compared to the first- and second-order models for adults and fledglings. In both groups, loglik becomes more positive from zeroth to first order and from first to second order, indicating that second-order structure is indeed present, and best explains both groups’ sequences (Figure 4). A greater improvement from zeroth to first order is seen in the adults than the fledglings, suggesting the presence of stronger first-order structure in adults. Adults then show a smaller improvement from first to second order, likely because their second-order structure is driven by strong adjacent-call structure (already captured by the first-order model) nested within their triplets of calls. In contrast, fledglings show comparable improvements from zeroth- to first-order models and first- to second-order models, suggesting their triplets of calls do not build as much on existing first order structure—likely due to the greater variability and flexibility present in fledgling sequences than in adults.

**Figure 4.**
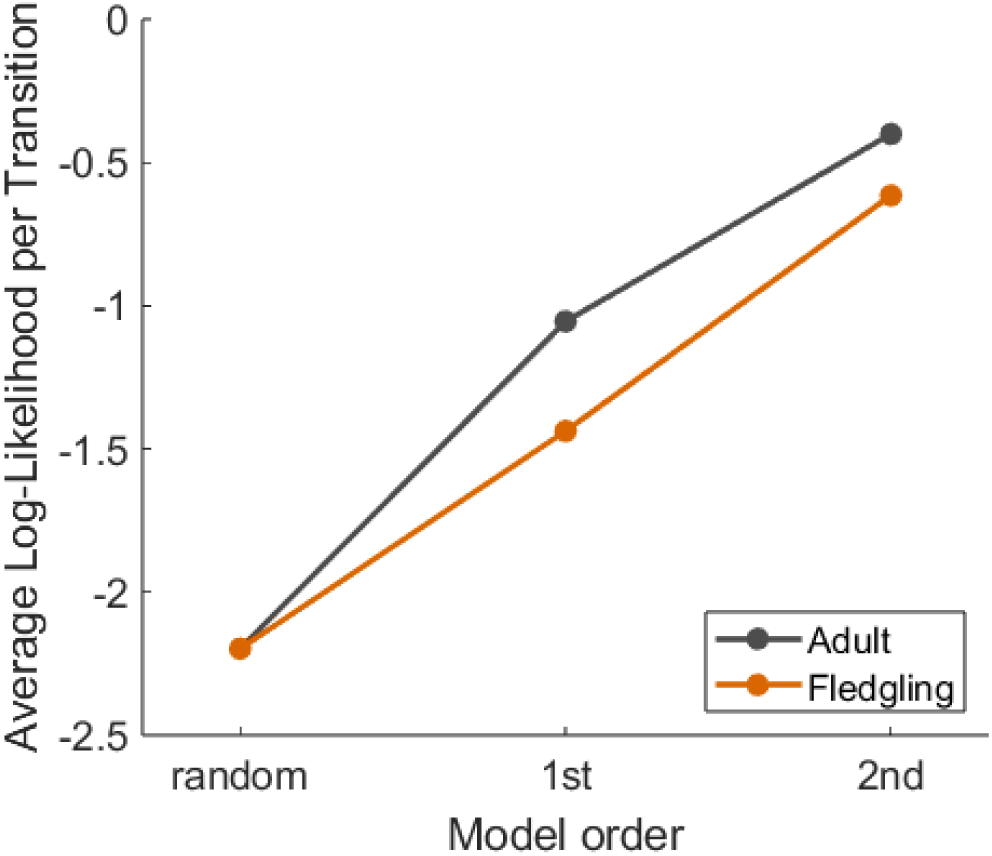
Average log-likelihoods per transition from zeroth order (random) to first- to second-order Markov models of adult and fledgling magpie vocal sequences. Trendlines are both averaged over additive smoothing constant *k* = 0.0001-1 (see Figures S.1. & S.2. in electronic supplementary material for full *k* comparison). While logliks are normalised over differences in dataset size between adults and fledglings, values should only be compared relatively, and not taken as absolutes, due to different baseline call frequencies, sequence lengths and transitions observed in each dataset.

## Discussion

Prior to this study, evidence of combinatorial structure in non-song repertoires beyond simple two-call combinations was scant and limited to primate studies (Bosshard et al., 2024; Girard-Buttoz, Zaccarella, et al., 2022). Building on earlier evidence of first-order structure within and between magpie calls (Walsh et al., 2023), we show that their extensive call sequences contain at least second-order structure. Specifically, we show that magpie sequences—which range from simple bigrams to 15-call sequences—are better explained by second-order than first-order Markov models (Figure 4), with overlapping significant triplets of calls suggestive of even further structure (Figure 5). The presence of ordering rules between segments within calls (Walsh et al., 2023), overlapping significant second-order transitions and recombination of calls and shorter sequences into longer ones, provides compelling evidence that magpie sequences exhibit hierarchical structure rather than simple adjacency chains. While work on common marmosets showed some of the first evidence of second-order structure and recombination at the semantic sequence level (Bosshard et al., 2024)—features that are consistent with hierarchy—magpies are the first species shown to exhibit ordered structure and recombination across two distinct levels, with second-order combinatorial rules operating over calls that are themselves formed from ordered segments (Walsh et al., 2023), providing stronger evidence of hierarchical organisation. We also capture the ontogeny of internal call sequence structure for the first time in any non-human animal, showing that fledglings display similar first-order but more variable second-order structure than adults—supporting a process whereby fledglings learn ordering rules during a period of flexible, exploratory sequencing.

**Figure 5.**
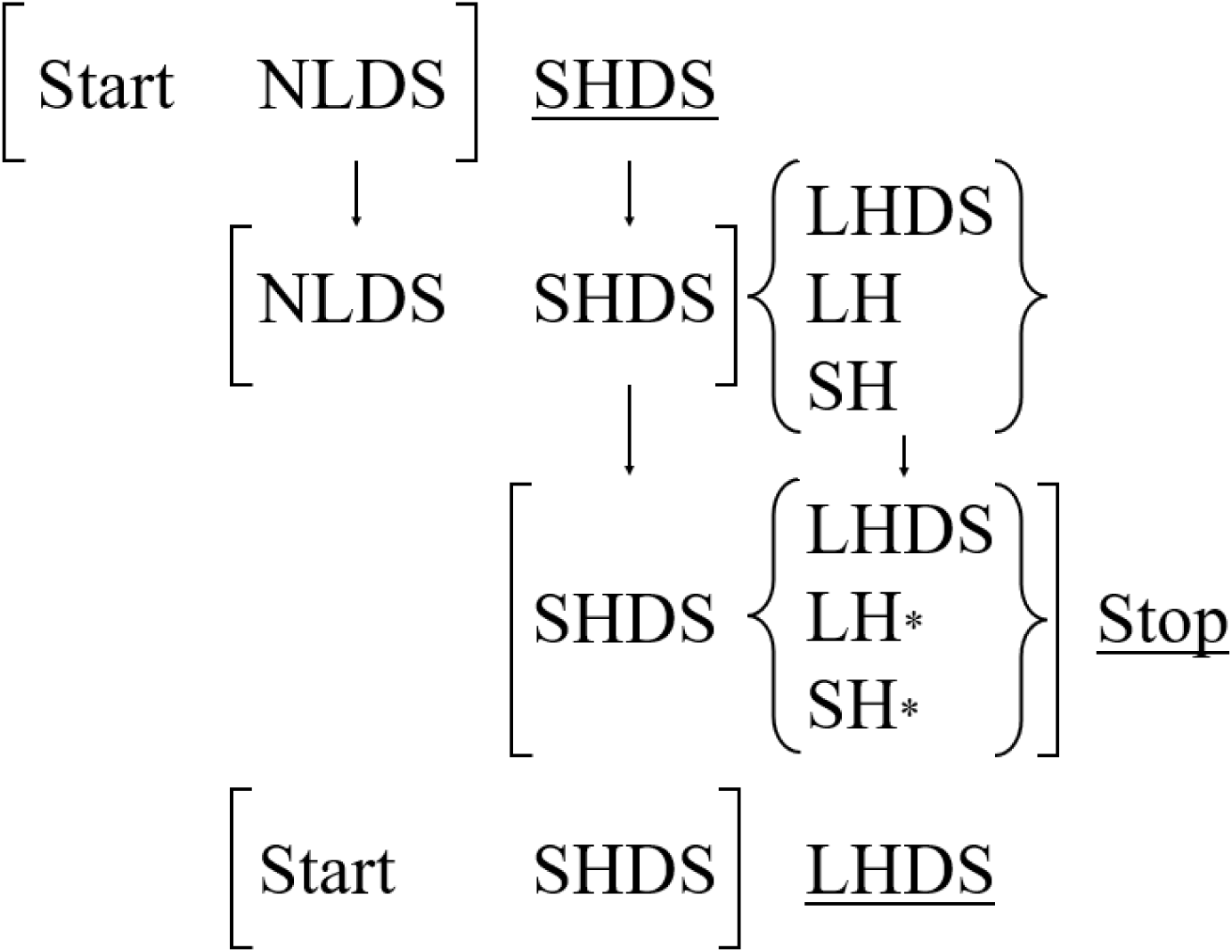
Overlapping second-order transitional dependencies within of two common sequences (SHDS-LHDS and NLDS-SHDS-LHDS produced 453 times and 356 times respectively in the dataset). Each square bracket represents a pair of calls or the Start state and the following call that significantly transition to a third call which is underlined—or in the case of several possible options—in curly brackets. ‘*’ indicates a transition that is highly probable (tP = 0.99) but insignificant (likely due to the low abundance of LH and SH in the dataset). The presence of transitional dependencies spanning up to five sequential positions within a sequence, paired with the recombination of segments within calls, calls within sequences and sequences within longer sequences (i.e. SHDS-LHDS in NLDS-SHDS-LHDS) suggests the presence of hierarchical structuring.

### Adult sequence structure

Magpie sequences show considerable flexibility, with adults producing at least 74 sequence types—many of which contain all four segments (*DS*, *NL, LH* and *SH*) present in a variety of single- and multi-segment calls (Walsh et al., 2023). We find that sequences exhibit predictable second-order dependencies, with significant transitions overlapping within longer, recombined sequences. For example, all the constituent triplets that make up the sequence *‘NLDS-SHDS-LHDS’* represent significant second-order transitions, indicating dependencies spanning up to five sequential positions (see Figure 5). *‘SHDS-LHDS’*, ‘*DS-SHDS’, ‘DS-SHDS-LHDS’* and *‘DS-SHDS-SH’*—all independently produced sequences—also contain significant, overlapping second-order transitions and together with ‘NLDS-SHDS-LHDS’, show how shorter sequences are recombined within a variety of longer sequences (Figure 3a). Although sequences were only modelled up to the second order, the presence of recombination and overlapping significant second-order transitions is suggestive of further structure—something that could be explored with additional data collection to boost sample sizes. Such analyses need not rely exclusively on Markov chain models (though see Morita et al., 2021 for example of 8^th^ order Markov structure in birdsong established using machine learning), but could instead benefit from more flexible frameworks drawn from birdsong research and computational linguistics (e.g. Hidden Markov Models, Katahira et al., 2011; Renewal Process Models, Kershenbaum et al., 2014; and Recurrent Neural Networks, Sainburg et al., 2019). Nevertheless, the presence of recombination at multiple levels—segments into calls, calls into sequences, and sequences into longer ones—using significant, overlapping second-order rules provides compelling evidence of hierarchical structuring.

In a semantic system, the presence of higher-order structure paired with recombination is suggestive of non-trivial compositionality. First, the presence of predictable ordering rules is, by itself, indicative of compositionality. While an alternative explanation for ordering rules could be morphological constraints (e.g., certain calls being physically easier or harder to produce after others, Raffelsiefen, 1999), every call type in the magpie repertoire occurs adjacent to every other at least once, ruling out production limitations. Given this, if order was not important for the meaning of a sequence, we would expect the probability of each transition to match chance levels based on the frequency of each call type in the dataset. Second, that calls are predicted by not just the current but also the previous state indicates that magpie sequences are not purely sequential—sequences are not reducible to pairwise dependencies, which is inconsistent with a simple additive system in which meaning accumulates call-by-call. Third, that independently produced sequences are reproduced within longer sequences with their internal structure preserved suggests that the combinatorial process occurring within the shorter sequences continues to operate once combined within the longer sequence. If meaning were associated with sequences holistically (i.e. non-compositionally) rather than built through structured, hierarchical ordering rules, there would be no expectation that internal structure be preserved in the constituent parts. The recombination of predictable sequences therefore suggests that structure is doing semantic work across levels, rather than longer sequences being stored and produced as chunks. These findings address key criticisms levelled at compositional research in Japanese tits (Suzuki et al., 2016) that highlighted the lack of flexible productivity (only one sequence was studied), and the lack of non-linear, hierarchical structure (Bolhuis et al., 2018; but see Suzuki et al., 2018 for rebuttal).

While the presence of such structure in a system known to be semantic is highly indicative of compositionality, identifying the individual meanings of further calls and sequences will allow us to confirm the presence of compositional meaning more concretely. The very flexibility and structural complexity that makes this system promising makes it is infeasible to experimentally determine the meaning to every individual call and sequence using playback methods used in single-sequence studies (Engesser et al., 2016; Suzuki et al., 2016; though see Mason, 2025 for evidence that magpie fledglings respond differently to a sequence and call). Berthet et al (2025) were able to identify differences in the meanings of a large number of bonobo calls and sequences by instead analysing the Euclidean distances of ‘Features of Circumstances’—a set of observationally collected variables including external events occurring at the time of the vocalisation, the behaviour of the signaller and the behavioural response of the receiver. While progress has been made in identifying probable meanings of some magpie calls and sequences from behavioural observation (see Walsh, 2023; Mason, 2025), this remains challenging due to the species’ mobility in flight—which makes one-to-one stimulus-response associations difficult to pinpoint—and the sheer diversity of the vocal repertoire. Adopting an approach like that used in bonobos—where meaning is identified from patterns across multiple contextual variables rather than from a single identified context—could enable broadscale analysis of the vast magpie repertoire.

### The development of sequence structure in fledglings

Previous findings suggest fledglings first combine innate segments into multi-segment calls, before learning to combine calls into complex sequences through social exposure (Mason, 2025; Mason et al., 2026). Here, we provide support for this theory: fledglings produce similar first-order structure to adults but more variable second-order structure, evident in their significant use of triplets of calls occurring at chance levels or entirely absent in adults. This mirrors song learning in birds, where flexible “subsong” precedes crystallization of adult song (Doupe & Kuhl, 1999; Thorpe, 1958), and early human language development, where children produce grammatically unconventional but communicatively effective utterances like “airplane all gone” (Braine, 1963; Winitz, 2020). While differences in adult and fledgling ordering rules could indicate age-specific sequences rather than heightened vocal flexibility during development, this is unlikely: while fledglings experience different social contexts (nutritional dependence, increased aggression), these are associated with discrete vocalizations (begging, grunting, discrete alarms) rather than sequences (Mason, 2025). Instead, the parallels between young magpies and humans discussed above offer compelling evidence for gradual acquisition of syntactic rules in a non-human species, underscoring the critical role of ontogenetic research in comparative linguistics.

Of the two ontogenetic studies of semantic call sequences in other species, both have focused on utterance length and diversity (Bortolato et al., 2023; Sigmundson et al., 2025), making this the first to investigate developmental emergence of ordering rules. Non-human primates show contrasting developmental modes: chimpanzee sequences mature over nearly a decade (Bortolato et al., 2023), while sooty mangabeys exhibit stable sequences from onset of production (Sigmundson et al., 2025). Magpies fall between these extremes: vocal building blocks develop rapidly—suggesting an innate basis (Mason, 2025; Mason et al., 2026)—while ordering rules emerge more slowly. Given that fledglings produce adult-length sequences by 200 days with no acoustic development in constituent units (Mason, 2025; Mason et al., 2026), their limitations likely reflect insufficient exposure to adult ordering rules rather than motor constraints.

While our previous research showed reduction of the repertoire following the period of heightened vocal flexibility occurs around the 200-day mark (Mason et al., 2026), it remains unclear whether this marks the emergence of adult-like second-order structure. In the present study it was not possible to examine developmental changes in greater detail beyond a broad comparison of adults and fledglings, due to sample size limitations. Finer-grained developmental analysis across additional breeding seasons would allow for pinpointing the timeline over which ordering rules stabilize. Moreover, a larger dataset would allow investigation of group-level differences in second-order structure. Given that magpie sequences are socially learned and differ substantially between groups (Walsh et al., 2024; Mason et al., 2026)—far exceeding the single example of a two-call greeting produced in opposite orders by two chimpanzee populations (Girard-Buttoz, Bortolato, et al., 2022)—second-order tendencies may also vary between groups. Notably, our previous work found that more sociable fledglings—those that spend more time with a greater number of social contacts—develop larger repertoires earlier (Mason et al., 2026). Investigating whether such fledglings also acquire stable and adult-like second-order structure sooner would offer novel insight into syntactic development in non-human animals and shed further light on the social drivers behind the evolution of syntax.

### Conclusion

The findings presented here not only deepen our understanding of vocal complexity in non-human animals but add to a growing body of evidence positioning non-primate species as valuable comparative models for exploring the mechanisms underpinning syntactic communication (Engesser et al., 2016, 2019, 2024; Suzuki et al., 2016, 2018, 2020; Walsh et al., 2023, 2024). The co-occurrence of syntactic communication and open-ended vocal production learning in this species challenges the dominant theory that syntax evolved solely to mitigate the constraints of genetically fixed repertoires (Bortolato et al., 2023; Nowak et al., 2000). We show here that young magpies learn the ordering rules with which to combine calls—a process more akin to usage learning—rather than the calls themselves, which previous work suggests are formed from innate building blocks (Mason, 2025; Mason et al., 2026). That a species capable of open-ended vocal production learning still appears to favour combining innate signals rather than learning new ones, suggests other constraints on communicative capacity—besides an inability to add new signals—may have driven the independent emergence of syntax in other lineages. This study underscores the value of both a whole-repertoire and comparative ontogenetic approach in revealing the mechanisms underlying syntactic communication and presents new opportunities for understanding how complex communication systems evolved.

