## Supplementary materials for "Socially learned call sequences reveal gradual development of higher-order structure"

| **Table S.1.** First-order Markov chain model matrix showing significant, Benjamini-Hochberg corrected *p* values from two-tailed permutation tests comparing first-order transition probabilities in magpie call sequences to chance expectation. First-order transitions are from a call or the *Start* state to a second call or the *Stop* state. Green cells indicate transitions that occurred significantly more than chance expectation and black cells, those that occurred significantly less than chance expectation when compared to a null distribution of 10,000 permutations of randomly generated sequences using the baseline frequency of calls and distribution of sequence lengths in the data. Adult values are given in bold text while fledgling values are italicised to aid in visual differentiation. ‘0.000’ indicates *p* < 0.0001. Blank cells indicate transitions that occurred at chance levels. | | | | | | | | | |
| --- | --- | --- | --- | --- | --- | --- | --- | --- | --- |
|  | Next call | DS | LH | LHDS | NL | NLDS | SH | SHDS | Stop |
| Starting call | Age |  |  |  |  |  |  |  |  |
| DS | Adult |  |  |  |  | **0.026** |  | **0.000** |  |
|  | Fledgling | *0.000* | *0.044* |  |  | *0.002* |  | *0.000* |  |
| LH | Adult |  |  |  |  | **0.007** |  | **0.010** | **0.000** |
|  | Fledgling |  |  |  |  | *0.002* |  | *0.005* | *0.000* |
| LHDS | Adult | **0.015** |  |  |  | **0.000** |  | **0.000** | **0.000** |
|  | Fledgling | *0.025* |  |  |  | *0.000* |  | *0.000* | *0.000* |
| NL | Adult |  |  |  |  |  |  |  | **0.008** |
|  | Fledgling |  |  |  |  | *0.006* |  | *0.003* | *0.025* |
| NLDS | Adult | **0.007** | **0.017** | **0.001** |  | **0.013** |  | **0.000** | **0.000** |
|  | Fledgling | *0.000* | *0.006* | *0.025* | *0.026* |  |  | *0.002* | *0.000* |
| SH | Adult |  |  |  |  |  |  |  | **0.000** |
|  | Fledgling |  |  |  |  | *0.025* |  |  | *0.000* |
| SHDS | Adult | **0.000** | **0.000** | **0.000** | **0.008** | **0.000** | **0.000** | **0.000** | **0.000** |
|  | Fledgling | *0.000* | *0.000* | *0.000* | *0.002* | *0.000* | *0.000* |  | *0.001* |
| Start | Adult |  | **0.007** | **0.000** |  | **0.000** |  |  |  |
|  | Fledgling |  | *0.001* | *0.000* |  | *0.000* | *0.012* | *0.000* |  |

| **Table S.2.** Second-order Markov chain model matrix showing significant, Benjamini-Hochberg corrected *p* values from two-tailed permutation tests comparing second-order transition probabilities in magpie call sequences to chance expectation. Second-order transitions are from a pair of calls or the *Start* state and next call to a third call or the *Stop* state. Green cells indicate transitions that occurred significantly more than chance expectation and black cells, those that occurred significantly less than chance expectation when compared to a null distribution of 10,000 permutations of randomly generated sequences using the baseline frequency of calls and distribution of sequence lengths in the data. Adult values are given in bold text while fledgling values are italicised to aid in visual differentiation. ‘0.000’ indicates *p* < 0.0001. Transitions that occurred significantly above chance but at a count ≤ 5 are indicated with asterisks. Blank cells indicate transitions that occurred at chance levels. The rows containing NA indicate that the starting pair of calls did not occur at all within the fledgling dataset. The final 13 rows of the table indicate starting pairs that did not occur at all within the adult dataset. | | | | | | | | | |
| --- | --- | --- | --- | --- | --- | --- | --- | --- | --- |
|  | Third call | DS | LH | LHDS | NL | NLDS | SH | SHDS | Stop |
| Starting pair | Age |  |  |  |  |  |  |  |  |
| DS-DS | Adult |  |  |  |  |  |  |  |  |
|  | Fledgling |  |  |  |  |  |  |  |  |
| DS-LH | Adult |  |  |  |  |  |  |  |  |
|  | Fledgling | *NA* | *NA* | *NA* | *NA* | *NA* | *NA* | *NA* | *NA* |
| DS-LHDS | Adult |  |  |  |  |  |  |  |  |
|  | Fledgling |  |  |  |  |  |  |  |  |
| DS-NL | Adult |  |  |  |  |  |  |  |  |
|  | Fledgling |  |  |  |  |  |  |  |  |
| DS-NLDS | Adult |  |  |  |  |  |  |  |  |
|  | Fledgling |  |  |  |  |  |  |  |  |
| DS-SH | Adult |  |  |  |  |  |  |  |  |
|  | Fledgling |  |  |  |  |  |  |  |  |
| DS-SHDS | Adult |  |  | **0.013** |  |  | **0.000** |  |  |
|  | Fledgling |  |  | *0.000* |  |  | *0.000* |  |  |
| LH-NL | Adult |  |  |  |  |  |  |  |  |
|  | Fledgling |  |  |  |  |  |  |  |  |
| LH-NLDS | Adult |  |  |  |  |  |  | **0.013*** |  |
|  | Fledgling |  |  |  |  |  |  |  |  |
| LHDS-NLDS | Adult |  |  |  |  |  |  | **0.000*** |  |
|  | Fledgling |  | *0.015** | *0.000** |  |  |  |  |  |
| NL-DS | Adult |  |  |  |  |  |  |  |  |
|  | Fledgling |  |  |  |  |  |  |  |  |
| NL-NLDS | Adult |  |  |  |  |  |  |  |  |
|  | Fledgling |  |  |  |  |  |  |  |  |
| NL-SHDS | Adult |  |  |  |  |  |  |  |  |
|  | Fledgling |  |  |  |  |  | *0.006** | *0.021** |  |
| NLDS-DS | Adult |  |  |  |  |  |  |  |  |
|  | Fledgling |  |  |  |  |  |  |  |  |
| NLDS-LH | Adult |  |  |  |  |  |  |  |  |
|  | Fledgling |  |  |  |  |  |  |  | *0.043** |
| NLDS-LHDS | Adult |  |  |  |  |  |  |  | **0.005*** |
|  | Fledgling |  |  |  |  |  |  |  | *0.000* |
| NLDS-NL | Adult |  |  |  |  |  |  |  | **0.038** |
|  | Fledgling |  |  |  |  |  |  |  | *0.011* |
| NLDS-NLDS | Adult |  |  |  |  |  |  | **0.000** | **0.000** |
|  | Fledgling |  |  |  |  |  |  | *0.000* | *0.000* |
| NLDS-SH | Adult |  |  |  |  |  |  |  |  |
|  | Fledgling |  |  |  |  |  |  |  |  |
|  | Third call | DS | LH | LHDS | NL | NLDS | SH | SHDS | Stop |
| Starting pair | Age |  |  |  |  |  |  |  |  |
| NLDS-SHDS | Adult |  | **0.005** | **0.000** |  | **0.038** | **0.016** |  | **0.000** |
|  | Fledgling |  | *0.000* | *0.000* |  | *0.003* |  |  |  |
| SH-LHDS | Adult |  |  |  |  |  |  |  |  |
|  | Fledgling | *NA* | *NA* | *NA* | *NA* | *NA* | *NA* | *NA* | *NA* |
| SH-NLDS | Adult |  |  |  |  |  |  |  |  |
|  | Fledgling | *0.014** |  |  |  |  |  |  |  |
| SHDS-DS | Adult | **0.013*** |  |  |  |  |  |  |  |
|  | Fledgling |  |  |  |  |  |  |  |  |
| SHDS-LH | Adult |  |  |  |  |  |  |  |  |
|  | Fledgling |  |  |  |  |  |  |  | *0.003* |
| SHDS-LHDS | Adult |  |  |  |  |  |  |  | **0.005** |
|  | Fledgling |  |  |  |  |  |  |  | *0.000* |
| SHDS-NLDS | Adult |  |  |  |  |  |  |  |  |
|  | Fledgling |  |  |  |  |  |  | *0.000** |  |
| SHDS-SH | Adult |  |  |  |  |  |  |  |  |
|  | Fledgling |  |  |  |  |  |  |  |  |
| SHDS-SHDS | Adult |  |  | **0.000** |  |  |  |  |  |
|  | Fledgling |  | *0.019* | *0.000* |  | *0.006* |  |  |  |
| Start-DS | Adult |  |  |  |  |  |  | **0.038** |  |
|  | Fledgling | *0.000* |  |  |  | *0.000* |  |  |  |
| Start-NL | Adult |  |  |  |  |  |  |  |  |
|  | Fledgling |  |  |  |  |  |  |  |  |
| Start-NLDS | Adult |  |  |  |  |  |  | **0.000** |  |
|  | Fledgling |  |  |  |  |  |  |  |  |
| Start-SH | Adult |  |  |  |  |  |  |  |  |
|  | Fledgling |  |  |  |  |  |  | *0.011** |  |
| Start-SHDS | Adult |  |  | **0.000** |  | **0.000** |  | **0.000** |  |
|  | Fledgling |  | *0.011* | *0.000* |  | *0.000* |  |  |  |
| LH-DS | Fledgling |  |  |  |  |  |  |  |  |
| LH-LH | Fledgling |  |  |  |  |  |  |  |  |
| LH-SHDS | Fledgling |  | *0.000** |  |  |  |  |  |  |
| LHDS-DS | Fledgling |  |  |  |  |  |  |  |  |
| LHDS-LH | Fledgling |  |  |  |  |  |  |  |  |
| LHDS-SHDS | Fledgling |  | *0.003** | *0.015** |  |  |  |  |  |
| NL-NL | Fledgling |  |  |  |  |  |  |  |  |
| NL-SH | Fledgling |  |  |  |  |  |  |  |  |
| SH-DS | Fledgling |  |  |  |  |  |  |  |  |
| SH-NL | Fledgling |  |  |  |  |  |  |  |  |
| SH-SHDS | Fledgling |  |  | *0.049** |  |  |  |  |  |
| SHDS-NL | Fledgling |  |  |  |  |  |  |  | *0.000** |
| Start-LHDS | Fledgling |  |  |  |  | *0.049* |  | *0.000** |  |

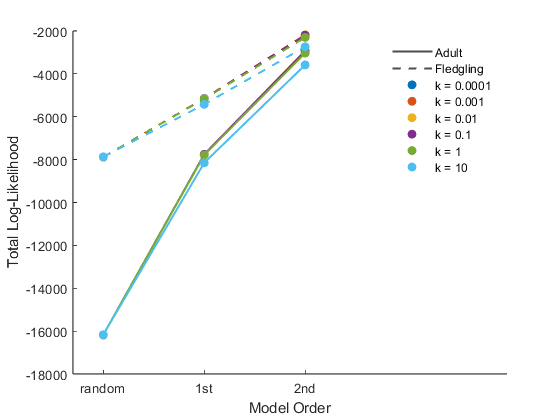

**Figure S.1.** Log-likelihoods per transition from zeroth- (random) to first- to second-order Markov models of adult and fledgling magpie vocal sequences. Each coloured trendline corresponds to a smoothing constant *k* = 0.0001-10. *k* = 10 caused underfitting as seen by its lower placement than the rest of the trendlines. As such only k = 0.0001-1 were used and averaged over in the analysis (Figure 4). While logliks are normalised over differences in dataset size between adults and fledglings, values should only be compared relatively, and not taken as absolutes, due to different baseline call frequencies, sequence lengths and transitions observed in each dataset.

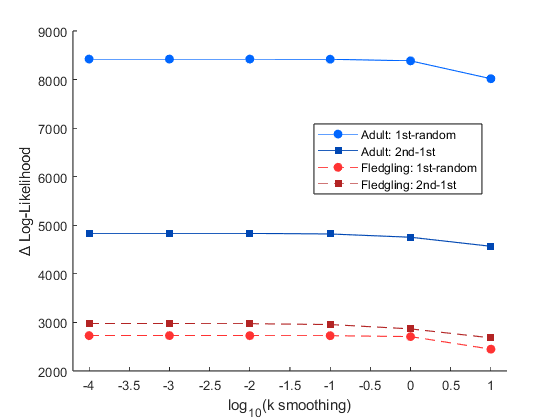

**Figure S.2.** Stability of smoothing constant k for each Markov model comparison—first-random and second-first order—for both adult and fledgling magpie call sequences. Values remain stable from k = 0.0001 (shown here on a logarithmic scale on the x axis as -4) to k = 1 (0 here), before dropping at k = 10 (1 here) due to over smoothing/underfitting. As such, k = 10 was excluded.
